# CCL5, CLEC11A, IL-7, IL-8, and IL-13: Diagnostic serum biomarkers of gastric cancer identified in a 48-multiplex panel

**DOI:** 10.1101/2025.08.15.670520

**Authors:** Jefim Brodkin, Tuomas Kaprio, Harri Mustonen, Alli Leppä, Arto Kokkola, Marko Salmi, Sirpa Jalkanen, Caj Haglund, Camilla Böckelman

## Abstract

**Background:** Gastric cancer is the fifth most common cancer worldwide and the fifth leading cause of cancer-related death. Its poor prognosis is primarily due to a late diagnosis and a lack of effective treatments for advanced disease.

**Methods:** We examined a patient cohort comprising 239 individuals who underwent surgery for histologically verified gastric adenocarcinoma in the Department of Surgery at Helsinki University Hospital between 2000 and 2009, comparing them to 48 healthy controls. We measured the serum protein concentrations for 48 different cytokines and growth factors using two of Bio-Rad’s premixed Bio-Plex Pro Human Cytokine 27-plex and 21-plex assay kits.

**Results:** Five serum biomarkers were identified as indicative of gastric cancer. Cancer patients had higher serum levels of CLEC11A [odds ratio (OR) 1.16, 95% confidence interval (CI) 1.08– 1.26, *p* = 0.004], IL-7 (OR 2.73, 95% CI 1.48–5.04, *p* = 0.014), IL-8 (OR 6.30, 95% CI 2.23–20.0, *p* = 0.017), and IL-13 (OR 2.67, 95% CI 1.33–5.37, *p* = 0.041). The CCL5 levels were lower in cancer patients compared with controls (OR 0.30, 95% CI 0.14–0.60, *p* = 0.014).

**Conclusions:** In a large cohort of 239 patients, we identified five biomarkers for which serum levels associated with gastric cancer: CCL5, CLEC11A, IL-7, IL-8, and IL-13. High serum levels of CLEC11A have not previously been associated with gastric cancer. Our results provide new support to further explore the effect of these inflammatory molecules and the role they play in gastric cancer. This may help identify novel noninvasive diagnostic methods as well as potential new druggable targets.

## Introduction

Gastric cancer (GC) is a common cancer worldwide. Although its incidence has decreased in the Western world, it remains one of the leading causes of cancer-related deaths. The five-year survival rate of GC in Finland from 2020 to 2022 was just above 30% [1], similar to other Western countries that do not screen for GC. Globally, however, GC has the fifth highest incidence and is the fifth most common cause of cancer-related death [2]. Its poor prognosis is mainly due to a late diagnosis and ineffective treatments for metastasized disease. To improve overall survival, new treatments and earlier diagnostics are needed. Diagnosis of GC is often delayed because the tumor might not initially cause any significant symptoms or the symptoms remain unspecific [3].

The diagnosis of GC relies on histological confirmation, obtained by endoscopy prior to any specific treatments. Imaging and blood samples also play a supportive role in diagnostics. Serum biomarkers, such as C-reactive protein (CRP), carcinoembryonic antigen (CEA), and carbohydrate antigen 19-9 (CA19-9), are used in the diagnosis and follow-up of GC patients [4, 5]. To date, well over 100 different proteins have been identified as diagnostics for GC [6]. For example, cancer antigens 12-5 (CA12-5) and 72-4 (CA72-4) can be used in diagnostics. Additionally, gastrin-17 and pepsinogen (PG) can be used for early GC detection [7]. However, they have not been widely implemented in clinical settings.

Liquid biopsies are a highly attractive area of research given that they provide a minimally invasive test producing individualized information. Cytokines and growth factors secreted by the tumor tissue—cancer cells as well as stromal cells—form the cancer secretome, which can also be examined from blood samples. Measuring the levels of proteins secreted by the tumor can yield indirect information about the tumor microenvironment (TME). In addition, serum biomarkers can include altered levels of normally expressed cytokines and growth factors, as well as tumor-originating microRNA or circulating tumor DNA (ctDNA), which are only found in patients with malignant disease.

In this study, we utilized Bio-Rad’s premixed Bio-Plex Pro Human Cytokine 27-plex and 21-plex assay kits to investigate possible serum biomarkers known to play a role in various cancers. We used logistic regression to identify statistically significant differences in serum levels comparing GC patients and healthy controls.

## Materials and methods

### Patients

The patient cohort comprised 239 individuals who underwent surgery for histologically verified gastric adenocarcinoma in the Department of Surgery at Helsinki University Hospital between 2000 and 2009. We excluded patients with a history of other malignant disease or synchronous cancer. In addition, we collected blood samples from 48 control patients with no known history of cancer. The median age of the patient cohort at the time of surgery was 65.6 years [interquartile range (IQR) 56.5–75.5; Supplementary Table 1]. Among patients, 117 (49.0%) were male, with a median survival of 2.34 years (IQR 0.90–9.95). For staging, we used the seventh version of the tumor-node-metastasis (TNM) classification [8]. In total, 49 (20.5%) patients had stage I cancer, 54 (22.6%) stage II, 94 (39.3%) stage III, and 42 (17.6%) stage IV. According to the Laurén classification, 84 (35.1%) patients had an intestinal histology, and 155 (64.9%) had diffuse or other types of cancer. Adjuvant chemotherapy was administered to 101 patients (44.3%) and 43 patients (19.2%) received adjuvant radiotherapy. Only 13 patients (5.4%) received neoadjuvant chemotherapy.

**Table 1.** Logistic regression analysis of biomarkers analyzed using the Bio-Rad’s premixed Bio-Plex Pro Human Cytokine 27-plex and 21-plex assays.

| | OR | 95% CI | $p$ value | FDR-corrected $p$ value |
| --- | --- | --- | --- | --- |
| **CCL5** | **0.30** | **0.14–0.60** | **0.001** | **0.014** |
| **CLEC11A** | **1.16** | **1.08–1.26** | **0.0001** | **0.004** |
| CTACK<sup>a</sup> | 1.37 | 0.36–4.74 | 0.626 | 0.776 |
| CXCL12<sup>a</sup> | 1.73 | 0.69–4.51 | 0.201 | 0.405 |
| Eotaxin<sup>a</sup> | 1.12 | 0.40–2.50 | 0.793 | 0.836 |
| FGF-Basic<sup>a</sup> | 2.52 | 0.74–9.00 | 0.118 | 0.334 |
| G-CSF<sup>a</sup> | 1.48 | 0.46–4.00 | 0.454 | 0.670 |
| GM-CSF<sup>a</sup> | 0.57 | 0.25–1.28 | 0.173 | 0.326 |
| GROα<sup>a</sup> | 0.85 | 0.33–1.93 | 0.714 | 0.790 |
| HGF<sup>a</sup> | 1.13 | 0.33–3.46 | 0.836 | 0.836 |
| IFN-α2<sup>a</sup> | 6.45 | 1.09–41.4 | **0.044** | 0.324 |
| IFN-γ<sup>a</sup> | 1.84 | 0.73–4.60 | 0.183 | 0.405 |
| IL-10<sup>a</sup> | 1.04 | 0.34–3.36 | 0.946 | 0.987 |
| IL-12p40<sup>a</sup> | 2.10 | 0.66–6.99 | 0.217 | 0.326 |
| IL-12p70<sup>a</sup> | 1.98 | 0.83–5.24 | 0.142 | 0.324 |
| **IL-13<sup>a</sup>** | **2.67** | **1.33–5.37** | **0.005** | **0.041** |
| IL-15<sup>a</sup> | 442 | 1.20–126 000 000 | 0.136 | 0.324 |
| IL-16<sup>a</sup> | 0.99 | 0.26–3.62 | 0.987 | 0.987 |
| IL-17<sup>a</sup> | 1.55 | 0.77–3.03 | 0.209 | 0.405 |
| IL-18<sup>a</sup> | 1.38 | 0.49–3.45 | 0.501 | 0.707 |
| IL-1RA<sup>a</sup> | 0.54 | 0.17–1.77 | 0.288 | 0.399 |
| IL-1α<sup>a</sup> | 1.33 | 0.77–2.34 | 0.316 | 0.515 |
| IL-1β<sup>a</sup> | 2.04 | 0.56–7.88 | 0.291 | 0.501 |
| IL-2<sup>a</sup> | 1.65 | 0.92–3.01 | 0.097 | 0.302 |
| IL-2RA<sup>a</sup> | 4.59 | 0.81–28.9 | 0.091 | 0.324 |
| IL-4<sup>a</sup> | 4.01 | 1.28–14.9 | **0.021** | 0.128 |
| IL-5<sup>a</sup> | 104 | 1.02–581 000 | 0.143 | 0.324 |
| IL-6<sup>a</sup> | 0.62 | 0.33–1.16 | 0.144 | 0.324 |
| **IL-7<sup>a</sup>** | **2.73** | **1.48–5.04** | **0.001** | **0.014** |
| **IL-8<sup>a</sup>** | **6.30** | **2.23–20.0** | **0.001** | **0.017** |
| IL-9<sup>a</sup> | 1.30 | 0.18–4.92 | 0.702 | 0.790 |
| IP-10<sup>a</sup> | 2.35 | 0.80–7.16 | 0.126 | 0.324 |
| LIF<sup>a</sup> | 0.72 | 0.18–2.81 | 0.643 | 0.724 |
| M-CSF<sup>a</sup> | 2.37 | 0.86–6.82 | 0.087 | 0.299 |
| MCP-1<sup>a</sup> | 1.14 | 0.34–3.33 | 0.815 | 0.836 |
| MCP-3<sup>a</sup> | 4.83 | 0.16–1 270 | 0.447 | 0.536 |
| MIF<sup>b</sup> | 1.93 | 0.96–4.14 | 0.076 | 0.293 |
| MIG<sup>a</sup> | 2.38 | 1.10–5.33 | **0.030** | 0.156 |
| MIP-1α<sup>a</sup> | 1.71 | 0.62–4.43 | 0.275 | 0.501 |
| MIP-1β<sup>a</sup> | 1.39 | 0.12–7.63 | 0.706 | 0.790 |
| PDGF-BB<sup>a</sup> | 0.69 | 0.17–2.46 | 0.591 | 0.763 |
| SCF<sup>a</sup> | 1.34 | 0.38–3.74 | 0.588 | 0.763 |
| TNF-α<sup>a</sup> | 2.32 | 0.68–8.32 | 0.161 | 0.393 |
| TNF-β<sup>a</sup> | 2.18 | 0.41–17.8 | 0.372 | 0.478 |
| TRAIL<sup>a</sup> | 2.41 | 0.95–7.76 | 0.073 | 0.293 |
| VEGF<sup>a</sup> | 2.14 | 0.65–7.53 | 0.214 | 0.326 |
| β-NGF<sup>a</sup> | 17.3 | 0.50–18 600 | 0.204 | 0.326 |

Abbreviations: OR, odds ratio; CI, confidence interval; FDR, false discovery rate.
<sup>a</sup>Statistical analyses were calculated using the logarithmic values of the biomarkers.
<sup>b</sup>Calculated as a dichotomous variable with the cutoff point at 800 pg/ml.

### Serum samples

Serum samples were obtained from patients after possible neoadjuvant therapy and shortly before surgery, aliquoted, and subsequently stored at –80°C until the multiplex assay was performed in 2018.

### Protein profiling

To determine the serum protein concentrations of cytokines and growth factors, we used two of Bio-Rad’s premixed Bio-Plex Pro Human Cytokine kits, the 27-plex assay (catalog no. M500KCAF0Y) and 21-plex assay (catalog no. MF0005KMII), on Bio-Rad’s Bio-Plex 200 system (Supplementary Table 2). Assays were performed according to the manufacturer’s instructions, however, only one-half of the recommended concentration levels for the number of detection antibodies, beads, and the streptavidin–phycoerythrin conjugate were used. We previously validated this approach [9, 10], which was successfully used in other cancer patient cohorts [11, 12].

**Table 2.** Associations between diagnostic biomarkers and clinicopathological variables among gastric cancer patients.

|  | CCL5<sup>high</sup><br>n = 100 (41.8%) | *p* value<sup>1</sup> | CLEC11A<sup>high</sup><br>n = 176 (73.9%) | *p* value<sup>1</sup> | IL-8<sup>high</sup><br>n = 146 (61.1%) | *p* value<sup>1</sup> | IL-13<sup>high</sup><br>n = 132 (55.2%) | *p* value<sup>1</sup> |
| --- | --- | --- | --- | --- | --- | --- | --- | --- |
| **Age** |  |  |  |  |  |  |  |  |
| < 66 years | 57 (46.3) | 0.152 | 91 (75.6) | 0.883 | 72 (58.5) | 0.405 | 69 (56.1) | 0.796 |
| ≥ 66 years | 43 (37.1) |  | 85 (73.3) |  | 74 (63.8) |  | 63 (54.3) |  |
| **Sex** |  |  |  |  |  |  |  |  |
| Male | 48 (41.0) | 0.896 | 94 (80.3) | **0.038** | 71 (60.7) | 0.900 | 67 (57.3) | 0.603 |
| Female | 52 (42.6) |  | 82 (67.8) |  | 75 (61.5) |  | 65 (53.3) |  |
| **Stage** |  |  |  |  |  |  |  |  |
| I | 12 (24.5) | 0.117<sup>2</sup> | 32 (65.3) | 0.222<sup>2</sup> | 21 (42.9) | **0.005<sup>2</sup>** | 26 (53.1) | 0.317<sup>2</sup> |
| II | 25 (46.3) |  | 42 (77.8) |  | 36 (66.7) |  | 25 (46.3) |  |
| III | 48 (51.1) |  | 69 (74.2) |  | 57 (60.6) |  | 57 (60.6) |  |
| IV | 15 (35.7) |  | 33 (78.6) |  | 32 (76.2) |  | 24 (57.1) |  |
| **Tumor invasion (pT)** |  |  |  |  |  |  |  |  |
| 1 | 9 (25.0) | **0.044<sup>2</sup>** | 25 (69.4) | 0.702<sup>2</sup> | 15 (41.7) | **0.007<sup>2</sup>** | 20 (55.6) | 0.377<sup>2</sup> |
| 2 | 14 (38.9) |  | 28 (77.8) |  | 19 (52.8) |  | 19 (52.8) |  |
| 3 | 38 (46.9) |  | 64 (80.0) |  | 55 (67.9) |  | 39 (48.1) |  |
| 4 | 39 (45.3) |  | 59 (68.6) |  | 57 (66.3) |  | 54 (62.8) |  |
| **Lymph node metastasis (pN)** |  |  |  |  |  |  |  |  |
| No | 26 (34.7) | 0.156 | 50 (66.7) | 0.113 | 40 (53.3) | 0.169 | 38 (50.7) | 0.484 |
| Yes | 70 (44.9) |  | 119 (67.4) |  | 98 (62.8) |  | 87 (55.8) |  |
| **Distant metastasis (M)** |  |  |  |  |  |  |  |  |
| No | 85 (43.1) | 0.395 | 143 (73.0) | 0.562 | 114 (57.9) | **0.027** | 108 (54.8) | 0.865 |
| Yes | 15 (35.7) |  | 33 (78.6) |  | 32 (76.2) |  | 24 (57.1) |  |
| **Laurén classification** |  |  |  |  |  |  |  |  |
| Intestinal | 39 (46.4) | 0.337 | 69 (82.1) | **0.044** | 60 (71.4) | **0.016** | 48 (57.1) | 0.685 |
| Diffuse and other | 61 (39.4) |  | 107 (69.5) |  | 86 (55.5) |  | 84 (54.2) |  |
| **MMR** |  |  |  |  |  |  |  |  |
| MMRp | 69 (40.1) | 0.477 | 126 (79.7) | 0.430 | 103 (59.9) | 0.236 | 96 (55.8) | 0.926 |
| MMRd | 19 (47.5) |  | 32 (80.0) |  | 28 (70.0) |  | 22 (55.0) |  |
| **EBV *ish*** |  |  |  |  |  |  |  |  |
| EBV negative | 84 (40.0) | **0.010** | 154 (73.7) | 0.464 | 128 (61.0) | 0.423 | 112 (53.3) | **0.009** |
| EBV positive | 7 (87.5) |  | 7 (87.5) |  | 6 (75.0) |  | 8 (100.0) |  |
| **p53 staining** |  |  |  |  |  |  |  |  |
| Aberrant | 70 (41.9) | 0.938 | 127 (76.5) | 0.258 | 102 (61.1) | 0.542 | 92 (55.1) | 0.978 |
| Wild type | 20 (42.6) |  | 32 (68.1) |  | 31 (66.0) |  | 26 (55.3) |  |

<sup>1</sup>Chi-square test. <sup>2</sup>Linear-by-linear test.
Abbreviations: CCL5, C–C motif chemokine 5; CLEC11A, C-type lectin domain family 11 member A;
IL, interleukin; MMRp/d, mismatch repair proficiency/deficiency; EBV *ish*, Epstein–Barr virus in situ hybridization.

### Tumor tissue microarray and immunohistochemistry

We also previously constructed a tumor tissue microarray, which we have immunostained for the following markers: MSI markers MSH2, MSH6, MLH1, and PMS2; p53; and EBER*ISH*. Our previous report provides a detailed description of our methods and the subsequent results [13].

## Statistical analysis

We used two-tailed *p* values and considered *p* < 0.05 as statistically significant. Statistical evaluations were calculated using IBM’s statistical software SPSS (IBM SPSS Statistics Version 28, International Business Machines Corp., NY, USA), R (R version 4.3.1, Foundation for Statistical Computing, Vienna, Austria), and GraphPad Prism (GraphPad Prism version 10.3.1 for MacOS, GraphPad Software, Boston, Massachusetts USA). Values falling below the standard curve were replaced by the minimum value obtained on the standard curve divided by the square-root of 2. This replacement was performed only if less than 10% of values did not fall on the standard curve, otherwise they were considered missing. We employed logistic regression to investigate the differences in the serum levels of the biomarkers between patients and healthy controls. The false discovery rate (FDR) for the multiple-test correction was applied [14]. We assessed associations between groups and continuous variables using the Mann–Whitney U-test and the Kruskal–Wallis test.

## Results

Of the 48 different cytokines and growth factors measured, we used logistic regression to identify differences in the serum levels between cancer patients and controls for eight biomarkers: CCL5, CLEC11A, interferon alpha 2 (IFN-*α*2), interleukin 4 (IL-4), IL-7, IL-8, IL-13, and monokine-induced by gamma interferon (MIG; Table 1). Of these biomarkers, IFN-*α*2, IL-4, and MIG were excluded given their non-significance after FDR correction. Cancer patients exhibited higher serum levels for CLEC11A [odds ratio (OR) 1.16, 95% confidence interval (CI) 1.08–1.26, *p* = 0.004], IL-7 (OR 2.73, 95% CI 1.48–5.04, *p* = 0.014), IL-8 (OR 6.30, 95% CI 2.23–20.0, *p* = 0.017), and IL-13 (OR 2.67, 95% CI 1.33–5.37, *p* = 0.041; Table 1). CCL5 levels were lower in cancer patients compared with controls (OR 0.30, 95% CI 0.14–0.60, *p* = 0.014; Table 1).

The five biomarkers were dichotomized using the maximum value of Youden’s index. The estimated cutoff point for CCL5 was 14 163 pg/ml, with a sensitivity of 70.8% and a specificity of 58.0%, differentiating GC patients from healthy controls (Figure 1a). The positive predictive value (PPV) was 90.8% and the negative predictive value (NPV) was 25.4%. The area under the curve (AUC) value was 0.661 (95% CI 0.571–0.751, *p* = 0.0004). The estimated cutoff point for CLEC11A was 129 585 pg/ml, with an AUC value of 0.691 (95% CI 0.610–0.773, *p* < 0.0001; Figure 1b). The estimated cutoff point for IL-7 was 11.28 pg/ml, with an AUC value of 0.624 (95% CI 0.533–0.714, *p* = 0.007; Figure 1c). The estimated cutoff point for IL-8 was 9.69 pg/ml, with an AUC value of 0.641 (95% CI 0.560–0.722, *p* = 0.002; Figure 1d). The estimated cutoff point for IL-13 was 1.49 pg/ml, with an AUC value of 0.592 (95% CI 0.505–0.680, *p* = 0.044; Figure 1e).

**Figure 1.**
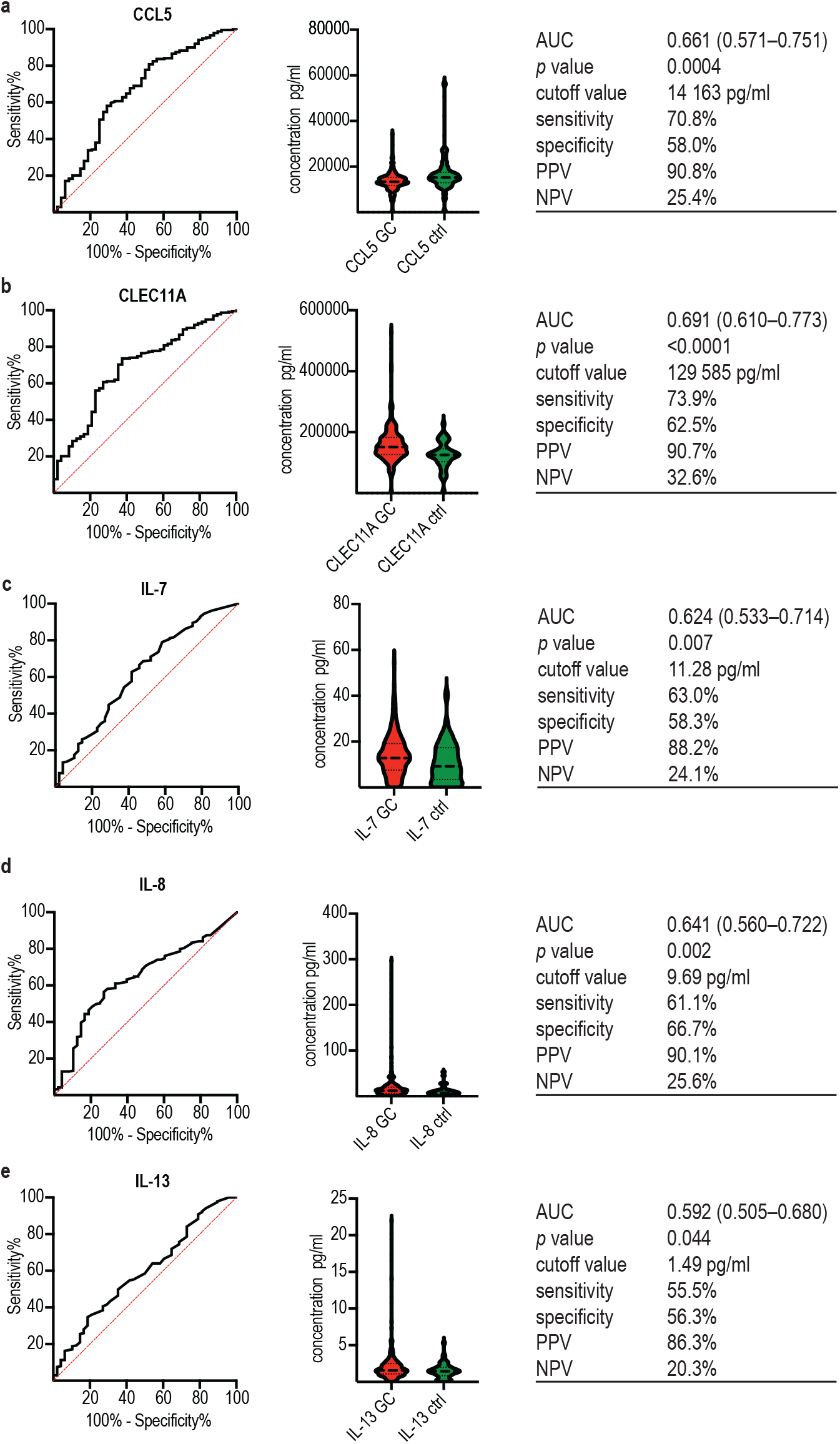
Receiver operating characteristic (ROC) curves and violin plots of the serum levels of a) CCL5, b) CLEC11A, c) IL-7, d) IL-8, and e) IL-13. Abbreviations: AUC, area under the curve; CCL5, C–C motif chemokine 5; CI, confidence interval; CLEC11A, C-type lectin domain family 11 member A; GC, gastric cancer; IL, interleukin; NPV, negative predictive value; PPV, positive predictive value; ROC, receiver operating characteristic.

### Associations between serum concentrations and clinicopathological variables

Higher serum levels of CCL5 were associated with Epstein–Barr virus (EBV) positivity (*p* = 0.010) and a higher pT value (*p* = 0.044; Table 2). Higher serum levels of CLEC11A were associated with male sex (*p* = 0.038) and an intestinal histology according to the Laurén classification (*p* = 0.044). A higher stage, pT value, and the occurrence of distant metastases as well as an intestinal histology based on the Laurén classification all associated with higher IL-8 serum concentrations. Higher serum levels for IL-13 associated with EBV positivity, with all EBV-positive patients in the cohort exhibiting high IL-13 serum levels (*p* = 0.009). IL-7 did not statistically associate with any of the variables we analyzed.

## Discussion

Logistic regression analysis revealed that among 48 analyzed biomarkers, CCL5, CLEC11A, IL-7, IL-8, and IL-13 served as diagnostic markers among GC patients. We found that these biomarkers can be used to differentiate GC patients from healthy controls. In a previous study, we used the same multiplex panel of 48 serum biomarkers to identify prognostic cytokines and growth factors in colorectal and pancreatic cancers [11, 12, 15]. However, the diagnostic biomarkers identified here were not prognostic in our patient population. Noninvasive methods for screening, diagnosing, and evaluating disease progression and aggressiveness remain under intense scrutiny. Blood samples and other easily obtained liquid biopsies represent an attractive area of research, given their relative speed, ease, and inexpensiveness as methods to implement in clinical use. Panels of six to nine selected serum markers have already been used to establish diagnostic marker combinations for GC [16, 17]. Previously, using a larger multiplex panel, Wu et al. [18] found changes in the levels of 11 different cytokines in GC. However, one limitation to their study was its rather small patient cohort of just 15 stage IA GC patients. Beyond cytokine panels, So et al. [19] developed a microRNA panel of 11 GC-associated microRNAs. Their findings, however, did not shed any further light on improving GC care. Preliminary results demand further validation, preferably in different patient cohorts, in order to provide justification for clinical use.

CCL5, together with its receptor C–C chemokine receptor type 5 (CCR5), is a well-known mediator of inflammation [20]. Furthermore, its role in various diseases associated with chronic inflammation remains under intense scrutiny. Many cancers, including gastric cancer, have been associated with increased CCL5–CCR5 signaling, which propagates cell proliferation, metastasizing, and angiogenesis [20]. Higher serum levels of CCL5 have thus been proposed as a diagnostic biomarker for GC [21, 22]. Another study found the serum levels of CCR5 (a CCL5 receptor) to be a diagnostic biomarker, but not CCL5 itself [23]. Contrary to some other studies, we found lower rather than higher levels of CCL5 were associated with GC. For instance, one study found an association between high pre-operative serum levels of CCL5 and occult peritoneal metastases [24]. We, however, observed no associations between CCL5 serum levels and the stage or other prognostic markers for GC. Serum CCL5 may originate from the tumor cells or from tumor stroma: fibroblasts, endothelial cells, and different immune cells. CCL5 secreted by tumor-associated macrophages [25] or CD4+ T-cells [26] have even been proposed as druggable targets. Our results related to lower serum levels for CCL5 associated with GC challenge previous findings. Taken together, the role of CCL5 is not as clear as previously thought, indicating the need for further exploration in larger patient cohorts.

CLEC11A was originally identified as a hematopoietic growth factor, but has also been associated with many metabolic functions as well as both blood cancers and cancers in the gastrointestinal tract [27]. Recently, CLEC11A was identified as also playing a significant role in GC-inducing cell migration and invasion as well as suppressing immunological activity in the tumor [28]. However, the diagnostic possibilities of CLEC11A have not previously been examined in a large patient cohort. Previous studies [28, 29] demonstrated that GC patients with a higher tumor expression of CLEC11A exhibited a worse prognosis. In another study, GC patients with increased CLEC11A gene expression levels had a better survival [30]. We found no prognostic effect of serum levels of CLEC11A, although we have not examined the tumor expression of CLEC11A. Furthermore, CLEC11A gene expression associated with the gene expressions of different immune cells [29]. The serum levels of CLEC11A have not been previously associated with cancer beyond one study, where it served as a predictive biomarker of chemotherapy in advanced hepatocellular cancer [31]. While CLEC11A is a novel diagnostic serum biomarker in GC, it seems to have a specific biological mechanism associated with the TME and immune response.

IL-7 is also a hematopoietic growth factor previously found to associate with gastrointestinal cancers like esophageal cancer, gastric cancer, and colorectal cancer [32], even though its biological mechanism in solid cancers is not precisely understood. In one study, the tumor expression of IL-7 was elevated in all these cancers compared with adjacent non-cancerous tissue [32]. However, the serum levels of IL-7 were elevated in colorectal and esophageal cancers, but not in GC. Our study, however, reveals that serum IL-7 can also be used to identify gastric cancer. Because the biological role of IL-7 in cancer is not well understood and has not been extensively studied, our results provide the rationale for further research.

IL-8, also known as C–X–C motif ligand 8 (CXCL8), is a pro-inflammatory cytokine that primarily attracts neutrophils, but which has been associated with various cancers [33]. Higher serum levels of IL-8 are already a known diagnostic indicator for gastric cancer, although they are not yet in clinical use [34–36]. The immunohistochemical expression of IL-8 was identified as higher in tumor tissue compared with adjacent normal tissue [37]. Furthermore, patients with a higher tissue expression of IL-8 have been shown to experience a worse prognosis [37]. Among gastric cancer patients, higher pretherapeutic serum levels of IL-8 have been linked to chemoresistance [38] and increased lymph node metastases [39]. Additionally, patients with high serum levels of IL-8 experience a shorter recurrence-free survival [39].

Accordingly, our results indicate associations with several disadvantageous properties, such as a higher stage of disease and distant metastases. The mechanistic effects of IL-8 have been extensively studied. The origins of serum IL-8 in GC have been connected to cancer-associated fibroblasts (CAF), further illustrating the close interaction between cancer cells and the TME [38, 40]. Beyond directly affecting cancer cells, IL-8 also works on T-cells by upregulating the PD-1 expression in them [41]. Additionally, IL-8 derived from CAFs or tumor-associated macrophages (TAM) induces PD-L1 expression in cancer cells [40, 42].

IL-13 is a well-known Th2-type cytokine regulating immune responses, which is also associated with the development of many malignant diseases [43]. In accordance with our results, high serum levels of IL-13 have been associated with gastric cancer as well as with pancreatic and esophageal cancers [44]. IL-13 has been identified as a necessary component in gastric metaplasia in a mouse model [45]. Our results did not reveal any further association with higher serum levels of IL-13 and clinical or pathological properties like the stage of disease or metastasis formation. Interestingly and similar to our results, a higher expression of IL-13 was previously reported as associated with EBV-infected B cells and Hodgkin’s lymphoma [46].

In our study, we used a rather large patient cohort of 239 patients composed almost entirely of gastric cancer patients undergoing surgery at Helsinki University Hospital in our selected time frame. Yet, a larger cohort is necessary for the further validation of our results. One benefit of an older patient cohort is that few patients received neoadjuvant treatment, allowing us to study cancer patients unaffected by chemotherapies or radiation therapy. Yet, these results need to be validated in neoadjuvant-treated patients given that chemotherapy changes the function of both cancer cells and the TME. Further research is also needed to examine whether these biomarkers are specific to only adenocarcinoma or whether they could also be altered in gastritis or other pre-malignant cases. In our study, we examined the effect of individual serum biomarkers instead of creating a model of several cytokines. The added statistical effect of numerous biomarkers is often greater; however, without a clear biological rationale for grouping them together, the effect may be merely statistical.

All five biomarkers we identified as diagnostically indicative of gastric cancer are chemokines and growth factors associated with inducing and sustaining inflammation. Most cases of GC are induced by chronic inflammation caused by *Helicobacter pylori* or EBV [47]. That said, *H. pylori* infection induces an immunosuppressive environment in the stomach through the recruitment of regulatory T-cells and anti-inflammatory macrophages already before cancer forms [47]. The role of different cytokines relating to interactions between cancer cells and the TME is under intense scrutiny, with further and more robust research needed [48]. Thus, our findings further underline the association between GC and inflammation. Ultimately, our results provide the rationale to further examine these cytokines and determine their biological role in cancer cells and the TME.

## Conclusions

In a large cohort of 239 patients using a multiplex panel screening for 48 serum markers, we identified five biomarkers whose serum levels associate with gastric cancer: CCL5, CLEC11A, IL-7, IL-8, and IL-13. High serum levels of CLEC11A have not previously been associated with gastric cancer. Our results provide new support for further exploring the effect of these inflammatory molecules and the role they play in gastric cancer. This may help identify novel noninvasive methods for diagnostics as well as new druggable targets.

## Acknowledgements

We thank Pia Saarinen and Maria Finne for their essential technical assistance, and Vanessa Fuller for English-language revisions.

## Funding

This study was financially supported by the Finnish Cancer Foundation (SJ, CH, and JB), Finska Läkaresällskapet (CH, CB, and JB), the Sigrid Jusélius Foundation (CH), the Emil Aaltonen Foundation (JB), the Finnish Medical Foundation (JB), Medicinska understödsföreningen Liv och Hälsa (CH, TK, and CB), the Waldemar von Frenckell Foundation (JB), the Paulo Foundation (JB), and the Orion Research Foundation (JB). The funders played no role in the study design, data collection and analysis, the decision to publish, or in producing the manuscript.

## Competing interests

The authors have no relevant financial or non-financial interests to disclose.

## Author contributions

Brodkin, Kaprio, Haglund, and Böckelman contributed to the study conception and design. Clinical data collection was performed by Leppä, Kokkola, Böckelman, and Haglund. Haglund collected the serum samples and Salmi and Jalkanen performed the cytokine measurements. Statistical analysis and interpretation were conducted by Brodkin, Kaprio, Mustonen, and Böckelman. The first draft of this manuscript was written by Brodkin, Kaprio, and Böckelman and all authors commented on the manuscript. All authors read and approved the final version of this manuscript.

## Data availability

The datasets supporting the conclusions of this article are included within the article and in its additional files. Other data used in this study are available from the corresponding author upon reasonable request.

## Ethics approval

Appropriate ethical permits were granted by the National Supervisory Authority of Health and Welfare and the Finnish Medicines Agency (permit for research conducted with human samples: Valvira Dnro 1004/06.01.03.01/2012), the hospital district of Helsinki and Uusimaa (permit number HUS/23/2024), and the Ethics Committee of Medicine of the Helsinki University Hospital (permit number HUS/1223/2021). Patient information, samples, and data were handled and stored in accordance with the Declaration of Helsinki and other local regulations.

## Consent to participate

Patients provided their written informed consent to participate in the study prior to providing blood samples.

## Abbreviations

AUC: area under the curve
CAF: cancer-associated fibroblast
CA12-5: cancer antigen 12-5
CA72-4: cancer antigen 72-4
CA19-9: carbohydrate antigen 19-9
CCL5: C–C motif chemokine ligand 5
CCL11: C–C motif chemokine ligand 11
CCR5: C–C chemokine receptor type 5
CEA: carcinoembryonic antigen
CI: confidence interval
CLEC11A: C-type lectin domain family 11 member A
CRP: C-reactive protein
ctDNA: circulating tumor DNA
CXCL8: C–X–C motif chemokine ligand 8
EBV: Epstein–Barr virus
FDR: false discovery rate
GC: gastric cancer
IL: interleukin
IQR: interquartile range
MMRd: mismatch repair deficiency
MMRp: mismatch repair proficiency
NPV: negative predictive value
PD-1: programmed cell death protein 1
PD-L1: programmed death ligand 1
PG: pepsinogen
PPV: positive predictive value
ROC: receiver operating characteristic
TAM: tumor-associated macrophage
TME: tumor microenvironment
TNM: tumor-node-metastasis classification

## Notes

### Competing Interest Statement

The authors have declared no competing interest.

