## additional file 1 for "CCL5, CLEC11A, IL-7, IL-8, and IL-13: Diagnostic serum biomarkers of gastric cancer identified in a 48-multiplex panel"

Additional File 1. Patient characteristics.

| | n = 239 (%) |
|---|---|
| **Age** | |
| < 66 years | 123 (51.5) |
| ≥ 66 years | 116 (48.5) |
| **Sex** | |
| Male | 117 (49.0) |
| Female | 122 (51.0) |
| **Stage** | |
| I | 49 (20.5) |
| II | 54 (22.6) |
| III | 94 (39.3) |
| IV | 42 (17.6) |
| **Tumor invasion (pT)** | |
| 1 | 36 (15.1) |
| 2 | 36 (15.1) |
| 3 | 81 (33.9) |
| 4 | 86 (36.0) |
| **Lymph node metastasis (pN)** | |
| No | 75 (32.5) |
| Yes | 156 (67.5) |
| **Distant metastasis (M)** | |
| No | 197 (82.4) |
| Yes | 42 (17.6) |
| **Laurén classification** | |
| Intestinal | 84 (35.1) |
| Diffuse and other | 155 (64.9) |
| **Adjuvant chemotherapy** | |
| No | 127 (55.7) |
| Yes | 101 (44.3) |
| **Adjuvant radiotherapy** | |
| No | 181 (80.8) |
| Yes | 43 (19.2) |
| **Neoadjuvant chemotherapy** | |
| No | 226 (94.6) |
| Yes | 13 (5.4) |
