## additional file 2 for "CCL5, CLEC11A, IL-7, IL-8, and IL-13: Diagnostic serum biomarkers of gastric cancer identified in a 48-multiplex panel"

Additonal file 2. Logistic regression of biomarkers analyzed using the Bio-Rad's premixed Bio-Plex Pro Human Cytokine 27-plex and 21-plex assays.

| | | **GC median** concentration (pg/ml) | **GC IQR** concentration (pg/ml) | **CTRL median** concentration (pg/ml) | **CTRL IQR** concentration (pg/ml) |
|---|---|---|---|---|---|
| CCL5 | C–C motif chemokine ligand 5 | 13 400 | 11 800–15 300 | 15 000 | 12 800–17 500 |
| CLEC11A | C-type lectin domain family 11 member A | 151 000 | 127 000–183 000 | 125 000 | 103 000–146 000 |
| CTACK | Cutaneous T cell-attracting chemokine | 1 340 | 977–1 870 | 1 340 | 981–1 920 |
| CXCL12 | C–X–C motif chemokine ligand 12 | 1 630 | 1 340–2 020 | 1 650 | 1 320–1 890 |
| Eotaxin | Eotaxin | 205 | 147–284 | 180 | 116–295 |
| FGF-Basic | Basic fibroblast growth factor | 37.8 | 33.9–41.9 | 34.2 | 32.3–38.2 |
| G-CSF | Granulocyte colony-stimulating factor | 207 | 162–258 | 208 | 156–256 |
| GM-CSF | Granulocyte-macrophage colony-stimulating factor | 2.32 | 1.91–4.52 | 2.77 | 2.07–4.68 |
| GRO$\alpha$ | Growth-regulated oncogene alpha | 265 | 241–293 | 277 | 254–292 |
| HGF | Hepatocyte growth factor | 811 | 621–1 050 | 770 | 632–1 020 |
| IFN-$\alpha$2 | Interferon alpha 2 | 6.86 | 4.97–8.91 | 6.31 | 4.57–8.07 |
| IFN- | Interferon gamma | 15.2 | 11.4–21.2 | 14.6 | 9.74–19.4 |
| IL-10 | Interleukin 10 | 7.41 | 5.64–13.7 | 5.85 | 5.59–6.11 |
| IL-12p40 | Interleukin 12 (p40) | 88.0 | 61.3–117 | 117 | 84.8–177.1 |
| IL-12p70 | Interleukin 12 (p70) | 4.56 | 2.00–8.78 | 3.44 | 1.42–8.81 |
| IL-13 | Interleukin 13 | 1.89 | 1.44–2.54 | 1.58 | 1.40–2.10 |
| IL-15 | Interleukin 15 | 176 | 110–290 | 61.9 | 59.7–69.4 |
| IL-16 | Interleukin 16 | 28.7 | 18.1–63.5 | 67.2 | 51.1–71.2 |
| IL-17 | Interleukin 17 | 8.57 | 6.71–10.5 | 8.57 | 6.85–10.4 |
| IL-18 | Interleukin 18 | 34.9 | 26.4–46.9 | 34.2 | 25.1–48.1 |
| IL-1RA | Interleukin 1 receptor antagonist | 179 | 135–228 | 192 | 134–273 |
| IL-1$\alpha$ | Interleukin 1 alpha | 7.90 | 4.13–12.0 | 7.90 | 5.55–8.35 |
| IL-1$\beta$ | Interleukin 1 beta | 1.34 | 1.19–1.69 | 1.38 | 1.14–1.72 |
| IL-2 | Interleukin 2 | 2.20 | 1.95–3.25 | 2.38 | 1.40–4.27 |
| IL-2RA | Interleukin 2 receptor antagonist | 74.4 | 59.0–96.1 | 63.8 | 52.0–85.5 |
| IL-4 | Interleukin 4 | 1.50 | 1.20–1.80 | 1.34 | 0.94–1.68 |
| IL-5 | Interleukin 5 | 21.7 | 18.5–76.2 | 3.76 | N/A |
| IL-6 | Interleukin 6 | 1.96 | 1.19–4.35 | 3.22 | 1.18–7.83 |
| IL-7 | Interleukin 7 | 15.3 | 12.2–20.7 | 15.1 | 11.6–19.6 |

Additonal file 2. Logistic regression of biomarkers analyzed using the Bio-Rad's premixed Bio-Plex Pro Human Cytokine 27-plex and 21-plex assays.

| | | | | | |
|---|---|---|---|---|---|
| IL-8 | Interleukin 8 | 14.7 | 9.36–22.1 | 9.14 | 6.02–13.2 |
| IL-9 | Interleukin 9 | 275 | 252–294 | 273 | 250–285 |
| IP-10 | Interferon gamma-induced protein 10 | 1 739 | 1 100–2 810 | 1590 | 939–2 430 |
| LIF | Leukemia inhibitory factor | 15.9 | 7.79–38.8 | 26.6 | 7.79–37.0 |
| M-CSF | Macrophage colony-stimulating factor | 27.3 | 22.7–36.2 | 25.1 | 18.8–32.6 |
| MCP-1 | Monocyte chemoattractant protein 1 | 33.8 | 26.2–45.2 | 29.4 | 24.9–49.4 |
| MCP-3 | Monocyte chemoattractant protein 3 | 1.37 | 1.37–4.27 | 1.37 | 1.37–2.12 |
| MIF | Macrophage migration inhibitory factor | 662 | 497–884 | 564 | 459–762 |
| MIG | Monokine induced by gamma interferon | 1 480 | 821–2 730 | 1 030 | 638–2 010 |
| MIP-1$\alpha$ | Macrophage inflammatory protein 1 alpha | 2.34 | 1.72–3.21 | 2.19 | 1.76–3.15 |
| MIP-1$\beta$ | Macrophage inflammatory protein 1 beta | 422 | 391–459 | 411 | 374–456 |
| PDGF-BB | Platelet-derived growth factor BB | 4 060 | 3 100–5 300 | 4 700 | 3 290–6 340 |
| SCF | Stem cell factor | 121 | 99.5–143 | 127 | 99.3–153 |
| TNF-$\alpha$ | Tumor necrosis factor alpha | 43.3 | 36.1–58.7 | 41.4 | 35.2–50.3 |
| TNF-$\beta$ | Tumor necrosis factor beta | 56.0 | 26.4–109 | 10.1 | 5.62–14.6 |
| TRAIL | TNF-related apoptosis-inducing ligand | 124 | 91.9–146 | 105 | 82.9–133 |
| VEGF | Vascular endothelial growth factor | 237 | 118–439 | 310 | 175–445 |
| $\beta$-NGF | Nerve growth factor beta | 2.50 | 1.79–5.37 | 1.39 | 0.78–2.00 |

Abbreviations: GC, gastric cancer; IQR, interquartile range
